# Advancing GABA-edited MRS Research through a Reconstruction Challenge

**DOI:** 10.1101/2023.09.21.557971

**Authors:** Rodrigo Pommot Berto, Hanna Bugler, Gabriel Dias, Mateus Oliveira, Lucas Ueda, Sergio Dertkigil, Paula D. P. Costa, Leticia Rittner, Julian P. Merkofer, Dennis M. J. van de Sande, Sina Amirrajab, Gerhard S. Drenthen, Mitko Veta, Jacobus F. A. Jansen, Marcel Breeuwer, Ruud J. G. van Sloun, Abdul Qayyum, Cristobal Rodero, Steven Niederer, Roberto Souza, Ashley D. Harris

**Author notes:** **Corresponding Author:** Hanna Bugler. Equal contribution (Rodrigo Pommot Berto, Hanna Bugler). co-senior authors (Roberto Souza, Ashley D. Harris).

## Abstract

**Purpose:** To create a benchmark for the comparison of machine learning-based Gamma-Aminobutyric Acid (GABA)-edited Magnetic Resonance Spectroscopy (MRS) reconstruction models using one quarter of the transients typically acquired during a complete scan.

**Methods:** The Edited-MRS reconstruction challenge had three tracks with the purpose of evaluating machine learning models trained to reconstruct simulated (Track 1), homogeneous *in vivo* (Track 2), and heterogeneous *in vivo* (Track 3) GABA-edited MRS data. Four quantitative metrics were used to evaluate the results: mean squared error (MSE), signal-to-noise ratio (SNR), linewidth, and a shape score metric that we proposed. Challenge participants were given three months to create, train and submit their models. Challenge organizers provided open access to a baseline U-NET model for initial comparison, as well as simulated data, *in vivo* data, and tutorials and guides for adding synthetic noise to the simulations.

**Results:** The most successful approach for Track 1 simulated data was a covariance matrix convolutional neural network model, while for Track 2 and Track 3 *in vivo* data, a vision transformer model operating on a spectrogram representation of the data achieved the most success. Deep learning (DL) based reconstructions with reduced transients achieved equivalent or better SNR, linewidth and fit error as conventional reconstructions with the full amount of transients. However, some DL models also showed the ability to optimize the linewidth and SNR values without actually improving overall spectral quality, pointing to the need for more robust metrics.

**Conclusion:** The edited-MRS reconstruction challenge showed that the top performing DL based edited-MRS reconstruction pipelines can obtain with a reduced number of transients equivalent metrics to conventional reconstruction pipelines using the full amount of transients. The proposed metric shape score was positively correlated with challenge track outcome indicating that it is well-suited to evaluate spectral quality.

## 1. INTRODUCTION

Magnetic resonance spectroscopy (MRS) is a non-invasive technique to quantify metabolite concentrations *in vivo*. One metabolite of particular interest to the neuroscience community is Gamma-Aminobutyric Acid (GABA), the primary inhibitory neurotransmitter(Paul G. Mullins et al., 2014). However, due to its low concentration and the chemical shift of its peaks, the GABA signal is overlapped by metabolites of more abundant concentrations, therefore it requires spectral editing for its quantification.

MEGA-PRESS(Harris et al., 2017; Paul G. Mullins et al., 2014) is the most commonly used technique to measure GABA. It consists of acquiring two subspectra. The edit-ON subspectra is acquired with an editing pulse at 1.9 ppm to modulate the GABA signal at 3 ppm while leaving other peaks at this frequency unaffected. The edit-OFF subspectra is acquired without applying an editing pulse. By subtracting the edit-OFF spectra from the edit-ON spectra, a difference spectra is obtained where the overlapping metabolites at 3 ppm are removed, revealing the 3 ppm GABA peak for quantification.

The subspectra subtraction affects edited-MRS SNR. Furthermore, edited-MRS is highly susceptible to motion artifacts and scanner instabilities resulting in subtraction artifacts(Ashley D. Harris et al., 2014). Therefore, GABA-edited MRS requires a large amount of measurements (*i.e.* transients), resulting in long scan times.

While preprocessing steps such as frequency and phase correction (FPC) and the removal of corrupted transients can improve GABA-edited MRS signal quality, they do not solve the trade-off between spectrum quality and scan time. To improve signal quality, we collect more data. However, longer scan times increase the probability of motion artifacts. Typical GABA-edited MEGA-PRESS acquisitions acquire on the order of 320 transients and take around 10 minutes per scan(A.L. Peek et al., 2023).

Machine learning (ML), more specifically deep learning (DL), has been proposed as an alternative preprocessing approach for conventional MRS(Chen et al., 2023; Kyathanahally et al., 2018; Lee & Kim, 2020; Wang et al., 2023) to reduce scan times and denoise spectra. However, DL approaches(Amirmohammad Shamaei et al., 2023; David J. Ma et al., 2022; Sofie Tapper et al., 2021) have yet to address reducing the long GABA-edited MRS scan times.

With an increase in machine learning-based solutions for MRS(Van De Sande et al., 2023), there is evidence to support that DL models can be used to reduce GABA-edited MRS acquisition times. For this challenge, reconstruction using one quarter of the typically acquired data was selected as the objective inspired by the results of magnetic resonance imaging (MRI) reconstruction challenges(Matthew J. Muckley et al., 2021; Youssef Beauferris et al., 2022) and the quality limitations of GABA-edited MRS data.

In the challenge, different models were compared to promote the investigation of DL techniques for processing edited-MRS data, with the ultimate goal of decreasing GABA-edited MRS acquisition times. In particular, participants of the challenge were asked to:

1. Create models which can reconstruct with high fidelity a GABA-edited MRS scan with only a quarter of the data traditionally acquired.
2. Assess the generalizability of their models from simulated data to *in vivo* data(Harris et al., 2023).

The results of the challenge were presented at the International Symposium on Biomedical Imaging (ISBI) conference in Cartagena, Colombia on April 18th, 2023. This publication includes a brief summary of model architectures and training parameters, metric results for each track, and a discussion on the difficulties encountered with model generalization and the use of *in vivo* specific metrics for DL model assessments. To make this benchmark reproducible, all the data and code pertaining to the challenge is available in its website (https://sites.google.com/view/edited-mrs-rec-challenge/home) and github repository (https://github.com/rmsouza01/Edited-MRS-challenge).

## 2. MATERIAL AND METHODS

### 2.1 Edited MRS Reconstruction Challenge Description

Participants were given over three months from December 16th, 2022 to March 27th, 2023 to prepare their solutions. The challenge was divided into three independent tracks; for Track 1, teams were provided a simulated ground truth development dataset and a noisy testing dataset and for Track 2 and Track 3, teams were provided a raw *in vivo* development dataset and testing dataset and were asked to submit the predictions from their methods on the provided testing datasets. The tracks were the following:

- **Track 1. Simulated Data:** Participants were provided with simulated ground truth data to which they could apply noise to create noisy transients to train and validate their models. These models were evaluated using noisy data generated by the challenge organizers and comparing each model’s 80 transient reconstruction to the associated simulated ground truths.
- **Track 2. Homogeneous *In Vivo* Data:** Participants were provided with an *in vivo* development dataset with 12 scans of 320 transients. Scans were collected with identical acquisition parameters to train and validate their models. These models were evaluated using a test set of 24 scans with near identical acquisition parameters as the development dataset and comparing each model’s 80 transient reconstructions to the 320 transient conventional reconstructions performed by GABA-edited MRS quantification software Gannet (Edden et al., 2014).
- **Track 3. Heterogeneous *In Vivo* Data:** Participants were provided with an *in vivo* development dataset with 30 scans comprising 320 transients, where scans were collected with different acquisition parameters (such as vendor, spectral points, and spectral width) to train and validate their models. These models were evaluated using a test set of 48 scans with similar parameters as the development dataset and comparing each model’s 80 transient reconstructions to 320 transient conventional reconstructions by Gannet(Edden et al., 2014).

While not required, teams were allowed and encouraged to use the data and model weights from the first tracks in the latter tracks, especially as starting points for training the models.

Model evaluation consisted of comparing four metrics of DL-based reconstructions: mean squared error (MSE), signal-to-noise ratio (SNR), linewidth, and shape score. In addition, a visual qualitative comparison was completed between the DL-based reconstructions and the 320 transient ‘target’ reconstructions. Each track was expected to increase in difficulty from simulated data to heterogeneous *in vivo* test data. While participants were not required to have one model across all three tracks, it was anticipated that teams would likely develop an initial architecture and fine tune it as tracks increased in difficulty.

### 2.2 Data

The challenge’s data includes both simulated and *in vivo* data divided into three tracks. Table 1 shows a summary of the datasets, with the size of each dataset and the source of the data. As the number of spectral points is an important parameter for the DL models, the data from Track 3 was balanced to have an equal number of acquisitions with 2048 and 4096 spectral points. This resulted in an imbalance in the number of acquisitions from each vendor, given the limitations of the available in vivo data.

**Table 1.** Summary of the datasets for the three tracks, with the source of the data, number of spectral points and number of development and testing samples. Track 3 has data from three different vendors and is split in four lines to show the amount of data from each vendor. Values in bold represent the total number of samples per track.

| Track # | Data Type | Data Source | Spectral points | Development samples | Testing samples |
| --- | --- | --- | --- | --- | --- |
| 1 | Simulated | FID-A matlab script | 2048 | **5,000**<br>ground truths | **1,000**<br>ground truths |
| 2 | *In Vivo* | Big GABA (GE) | 2048 | **12**<br>scans of 320<br>transients | **24**<br>scans of 80<br>transients |
| 3 | *In Vivo* | Big GABA (GE) | 4096 | 6<br>scans of 320<br>transients | 6<br>scans of 80<br>transients |
|  |  | Big GABA (Phillips) | 2048 | 12<br>scans of 320<br>transients | 24<br>scans of 80<br>transients |
|  |  | Big GABA (Siemens) | 4096 | 12<br>scans of 320<br>transients | 18<br>scans of 80<br>transients |
|  |  | **Total** |  | **30**<br>scans of 320<br>transients | **48**<br>scans of 80<br>transients |

#### 2.2.1 Simulated Data

The simulated data used in the challenge can be divided into two categories: ground-truth spectra and noisy transients. The ground truth data consists of noiseless free induction decays (FIDs) generated using FID-A(Robin Simpson et al., 2017), a Matlab simulation and processing tool for MRS. FIDs were generated with the resonance of 22 metabolites (alanine, ascorbate, aspartate, beta-hydroxybutyrate, Cr, GABA, glutamine, glutamate, glutathione, glycine, glycerophosphocholine, glucose, myo-inositol, lactate, N-acetylaspartate, N-acetylaspartateglutamate, phosphocholine, phosphocreatine, phosphoethanolamine, scyllo-inositol, serine, taurine), five macromolecules (MM 09, 12, 14, 17, 20) and one lipid (Lip20)(Jamie Near et al., 2013). Each component’s concentration was sampled independently from a normal distribution with the mean values reported from the literature(Jamie Near et al., 2013) and +/- 10% standard deviation. The acquisition parameters were: magnetic field strength of 3T, MEGA-PRESS variant (FID-A’s MegaPressShaped_fast(Yan Zhang et al., 2017)), edit-ON pulse at 1.9 ppm for 14 ms, edit-OFF pulse at 7.46 ppm for 14 ms, editing pulses interleaved between ON and OFF each TR, TR/TE= 2 s/68 ms, spectral width of 2 kHz, and 2048 points.

To generate the noisy transients, noise was added to the ground truth FIDs using an in-house python script provided to participants to mimic the artifacts found in *in vivo* spectra. The added noise consisted of a random combination of time domain Gaussian distributed amplitude noise, frequency shifts (variations along the frequency axis), and phase shifts (variations along the real-complex plane). Noise was added to each FID as follows:

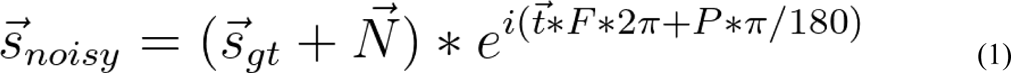

Where *S_gt_* is the ground truth FID, *N* is the amplitude noise, *t* is the time array for the acquisition, *F* is the frequency noise and *P* is the phase noise. The noises are obtained by sampling random normal distributions with flexible standard deviation that can be adjusted by teams for the desired noise level.

For Track 1, 5,000 GABA-edited simulated ground truth FID pairs (i.e. 5,000 edit-ON and 5,000 edit-OFF) were provided for the development dataset, to which participants were expected to create noisy transients using the in-house python scripts or any other method developed by the teams. In contrast, the testing dataset consisted of 1,000 GABA-edited simulated scans of 80 noisy transients (40 edit-ON and 40 edit-OFF), where noise was applied to the ground truth simulated spectra by the challenge organizers. The ground-truth difference spectra formed the target for the reconstructions.

#### 2.2.2 *In Vivo* Data

The *in vivo* data used in the challenge is from the Big GABA(Mark Mikkelsen et al., 2017) public dataset and FID-A(Robin Simpson et al., 2017) was used to extract the FIDs from the raw data files. For Track 2, *in vivo* scans of near identical (homogeneous) parameters were used as an intermediary step towards broader generalization as seen in track 3. The parameters associated with the data from Track 2 were the following: magnetic field strength of 3T, General Electric (vendor), CHESS water suppression, with 15 ms editing pulses at 1.9 ppm (ON) and

7.46 ppm (OFF) interleaved between ON and OFF each TR, TR/TE= 2 s/68 ms, spectral width of 2 kHz, 2048 points, 8 step phase cycling, and 320 transients. The development dataset consisted of 12 scans of 320 transients provided as raw FIDs each scan with its own associated target reconstruction by Gannet using the full 320 transients. The testing dataset consisted of 24 scans where the first 80 transients acquired in the scan were provided to participants and subsequently compared by the organizers against the target 320 transient reconstructions performed by Gannet.

For Track 3, the acquisition parameters were a combination of the following: magnetic field strength of 3T, GE, Siemens, or Philips (vendor), CHESS or MOIST or WET water suppression, with 15 ms editing pulses (GE and Philips) at on 1.9 ppm (ON) and 7.46 ppm (OFF) interleaved between ON and OFF each TR, TR/TE= 2 s/68 ms, spectral width of 2 kHz, 4 kHz, or 5 kHz, 2048 or 4096 points, 8 or 16 step phase cycling, and 320 transients. For Track 3, the development dataset consisted of 30 scans of 320 transients provided as raw FIDs each scan with its own associate target reconstruction by Gannet using the full 320 transients. The testing dataset consisted of 48 scans where the first 80 transients acquired in the scan were provided to participants. Results were compared by the organizers against the target, which was the full 320 transient reconstruction performed by Gannet.

### 2.3 Metrics

Four metrics were used to evaluate the reconstructions. They were chosen as a combination of traditional DL and MRS metrics and a new proposed metric. The choice of metrics was meant to evaluate different aspects of the reconstructions.

#### 2.3.1 Mean Square Error (MSE)

The MSE is a traditional metric used for assessing similarity to a reference. The reference measurement used for Track 1 was the ground truth and for Track 2 and Track 3, the 320 transient ‘target’ reconstruction was used. MSE was measured between 2.5 ppm and 4 ppm. To ensure fair evaluation, a min-max normalization was applied between 2.5 ppm and 4 ppm on both the reference and output spectra. The normalization applied can be defined by the following equation:

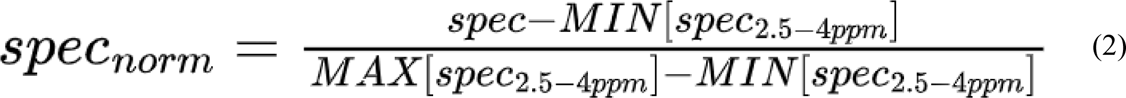

#### 2.3.2 Signal to Noise Ratio (SNR)

SNR is a traditional metric used to assess MRS data quality(Alexander Lin et al., 2021; In-Young Choi et al., 2021; Jamie Near et al., 2021). As edited-GABA is the signal of interest, GABA SNR as per Eq. 3 was used.

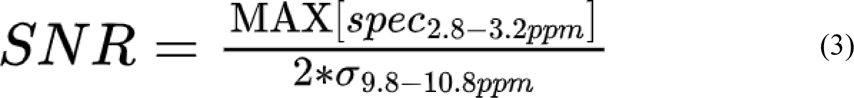

Where the numerator is the maximum value of the GABA peak found between 2.8 ppm and 3.2 ppm, and σ is the standard deviation of the signal between 9.8 ppm and 10.8 ppm after second order polynomial fitting of the region to avoid residual influences of other peaks and wandering baseline(Mark Mikkelsen et al., 2017). Similarly to MSE, to ensure the metric is not affected by scaling or offsets, a normalization is applied to maximize the GABA peak at 1 and the median of the spectra at 0. A different normalization from the one used for MSE was necessary to approximate the upfield mean value to zero and avoid deviating the metric due to GABA peak offsets. The normalization utilized can be defined by the following equation:

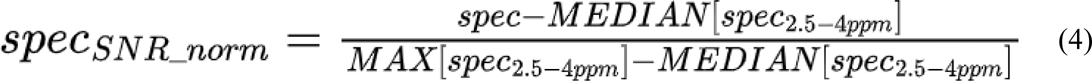

#### 2.3.3 Linewidth

Linewidth is also a traditional metric used to assess spectral quality(Alexander Lin et al., 2021; In-Young Choi et al., 2021; Jamie Near et al., 2021). Similarly to SNR, as GABA is the metabolite of focus for the challenge, its linewidth (full width at half maximum (FWHM)) was compared.

#### 2.3.4 Shape Score

In MRS, often visualization of spectra is an opportunity for qualitative assessments of the data overall and more specifically, the peak(s) of interest. As an alternative, the organizers proposed the shape score as a quantitative approach to replace the qualitative assessment through visual inspection. Similarly to MSE, the shape score focuses on the shape of the GABA and Glx peaks by comparing them to the expected result. This metric consists of calculating the correlation between the normalized ground truth simulation (Track 1) or target *in vivo* full 320 transient reconstruction (Track 2 and Track 3) and the normalized model reconstruction. For the GABA peak, a spectral window between 2.8 ppm and 3.2 ppm was used and for the Glx peak, a window between 3.55 ppm and 3.9 ppm was used (Equation 4). The shape score is calculated as a weighted sum equivalent to 60% of the GABA correlation and 40% of the Glx correlation. The weights were chosen to reflect the objective of the challenge, high quality GABA reconstruction, while ensuring reconstructions were considerate of other important peaks that contribute to model reliability. The formula for shape score is the following:

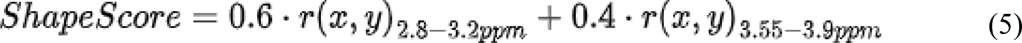

Where r is the Pearson’s correlation coefficient for the subscripted region, *x* is the model reconstruction and *y* is the target reconstruction. The 2.8 - 3.2 ppm region refers to the GABA peak and the 3.55 - 3.9 ppm region refers to the Glx peak.

#### 2.3.5 Fit Error

Challenge organizers opted to introduce the results of an additional metric, fit error, which was not used in the challenge’s ranking. As this metric is calculated based on an individual spectrum, two supplementary results are added for comparison: reference - which is the spectrum reconstructed by Gannet with the full 320 transients (*i.e.* the same spectrum used for the target previously); and control - which is the spectrum reconstructed by Gannet with only 80 transients

### 2.4 Models

In order to establish the challenge baseline and encourage participants to create models exceeding this starting point, the organizers provided the code and metric results for a simple U-NET model(Ronneberger et al., 2015). In this model, the 80 FIDs were converted to 40 difference spectra by an inverse Fourier Transform followed by subtraction, and the difference spectra was concatenated to form a 2D 2048×40 image. This image is passed through the 10 convolutional blocks of the U-Net and then averaged over the transient dimension at the end, resulting in a single 2048 elements array. Prior to presentation of the results at the 2023 ISBI conference, the challenge obtained submissions from three groups of researchers for each of the three tracks.

#### 2.4.1 Team Deep Spectral Divers: Spectrogram Vision Transformer

Team Deep Spectral Divers submitted a vision transformer combined with a multilayer perceptron decoder for all three tracks. The input to the model is the max-normalized zero-padded mean difference spectrogram obtained using 40 transients from ON and OFF FIDs converted via a short-time Fourier transform (STFT). Each track had its own model which was trained by initializing its weights to those of the previous track (*i.e.* track 2 model began training with track 1 final model weights). In addition, track 3 model had two sub-models which differed in the number of input spectral points where the 4096 spectral point sub-model was trained based on the final weights of the 2048 spectral point sub-model.

#### 2.4.2 Team Spectralligence: Covariance Matrix Convolutional Neural Network

Team Spectralligence submitted a convolutional neural network (CNN) whose architecture of the final layers varied per track(Julian P. Merkofer et al., 2023). The input to the model is a covariance matrix of transient measurements (spectral point values) which can operate arbitrarily on any number of transients. Track 1 model’s CNN configuration was followed by a flattening layer and subsequently a set of fully connected layers for model prediction. In contrast, track 2 and track 3 model CNN configurations were followed by an inverse fast Fourier transform (IFFT) operation and a recurrent neural network (RNN) to more carefully extract local features while avoiding overfitting.

#### 2.4.3 Team Dolphins: Spectral Image U-Net

Team Dolphins submitted a U-NET model with depth-wise channel attention module (DCAM). The DCAM block is introduced in each encoder block after the convolutional layer. This replaces the fully connected layers in the channel attention module with a 1D convolution layer after channel-wise global average pooling without dimensionality reduction. The input to the model is a 2D representation of concatenated spectra and the model was optimized using both the MSE and Dynamic Huber Loss. The model trained for track 1 was used as a starting point for training track 2 and track 3 where the parameters of the feature extraction layers are frozen and the predictive layers are fine-tuned to data of the new or shifted domain.

Further information regarding architecture or training of individual models can be obtained through open-access code source repositories in the appendices of this report.

## 3. RESULTS

The average metrics results for each track of the challenge are shown in Table 2.

**Table 2.** Metric values (mean +/- std) calculated for each team based on the procedures for evaluation of models on the test set for each track of the edited-MRS reconstruction challenge. Results in bold indicate the top result for a particular metric for a particular track and models with an asterisk indicate a result not considered in the final ranking.

| Section 1: Track 1 - Simulated Data |  |  |  |  |
| --- | --- | --- | --- | --- |
| Model | Mean Squared Error (MSE) | Signal-to-Noise Ratio (SNR) | Linewidth | Shape Score |
| Baseline U-Net* | 0.03452 ± 0.00748 | 91.3 ± 64.2 | 0.076661 ± 0.006877 | 0.7000 ± 0.1359 |
| Deep Spectral Divers | 0.00100 ± 0.00078 | 80.2 ± 21.8 | **0.076447 ± 0.000242** | 0.9994 ± 0.0004 |
| Spectralligence | **0.00089 ± 0.00077** | **2361.5 ± 768.2** | 0.076455 ± 0.000000 | **0.9996 ± 0.0002** |
| Dolphins | 0.08971 ± 0.02920 | 2.0 ± 0.6 | 0.343908 ± 0.547723 | 0.2366 ± 0.2620 |
| Section 2: Track 2 - Homogeneous *in vivo* Data |  |  |  |  |
| Model | Mean Squared Error (MSE) | Signal-to-Noise Ratio (SNR) | Linewidth | Shape Score |
| Baseline U-Net* | 0.03486 ± 0.03254 | 45.2 ± 24.4 | 0.147175 ± 0.042553 | 0.8283 ± 0.2013 |
| **Deep Spectral Divers** | 0.01529 ± 0.02008 | **48.9 ± 2.6** | **0.153865 ± 0.002583** | **0.9809 ± 0.0202** |
| Spectralligence | **0.00999 ± 0.01050** | 47.1 ± 23.1 | 0.163103 ± 0.006637 | 0.9769 ± 0.0233 |
| Dolphins | 0.03574 ± 0.01616 | 6.4 ± 1.6 | 0.171704 ± 0.044005 | 0.7037 ± 0.1312 |
| **Section 3: Track 3 - Heterogeneous *in vivo* Data** |  |  |  |  |
| **Model** | **Mean Squared Error (MSE)** | **Signal-to-Noise Ratio (SNR)** | **Linewidth** | **Shape Score** |
| Baseline U-Net* | 0.02420 ± 0.01711 | 35.6 ± 17.5 | 0.172427 ± 0.046778 | 0.8253 ± 0.1040 |
| **Deep Spectral Divers** | **0.01271 ± 0.03592** | **44.2 ± 10.6** | 0.182649 ± 0.030305 | **0.9486 ± 0.2090** |
| Spectralligence | 0.01697 ± 0.02586 | 33.9 ± 13.9 | **0.166503 ± 0.021886** | 0.9195 ± 0.2118 |
| Dolphins | 0.07279 ± 0.05466 | 5.9 ± 2.0 | 0.191105 ± 0.040382 | 0.3700 ± 0.4909 |

| Section 1: Track 1 - Simulated Data |  |  |  |  |
| --- | --- | --- | --- | --- |
| Model | Mean Squared Error (MSE) | Signal-to-Noise Ratio (SNR) | Linewidth | Shape Score |
| Baseline U-Net* | $0.03452 \pm 0.00748$ | $91.3 \pm 64.2$ | $0.076661 \pm 0.006877$ | $0.7000 \pm 0.1359$ |
| Deep Spectral Divers | $0.00100 \pm 0.00078$ | $80.2 \pm 21.8$ | **$0.076447 \pm 0.000242$** | $0.9994 \pm 0.0004$ |
| Spectralligence | **$0.00089 \pm 0.00077$** | **$2361.5 \pm 768.2$** | $0.076455 \pm 0.000000$ | **$0.9996 \pm 0.0002$** |
| Dolphins | $0.08971 \pm 0.02920$ | $2.0 \pm 0.6$ | $0.343908 \pm 0.547723$ | $0.2366 \pm 0.2620$ |
| Section 2: Track 2 - Homogeneous *in vivo* Data |  |  |  |  |
| Model | Mean Squared Error (MSE) | Signal-to-Noise Ratio (SNR) | Linewidth | Shape Score |
| Baseline U-Net* | $0.03486 \pm 0.03254$ | $45.2 \pm 24.4$ | $0.147175 \pm 0.042553$ | $0.8283 \pm 0.2013$ |
| Deep Spectral Divers | $0.01529 \pm 0.02008$ | **$48.9 \pm 2.6$** | **$0.153865 \pm 0.002583$** | **$0.9809 \pm 0.0202$** |
| Spectralligence | **$0.00999 \pm 0.01050$** | $47.1 \pm 23.1$ | $0.163103 \pm 0.006637$ | $0.9769 \pm 0.0233$ |
| Dolphins | $0.03574 \pm 0.01616$ | $6.4 \pm 1.6$ | $0.171704 \pm 0.044005$ | $0.7037 \pm 0.1312$ |
| Section 3: Track 3 - Heterogeneous *in vivo* Data |  |  |  |  |
| Model | Mean Squared Error (MSE) | Signal-to-Noise Ratio (SNR) | Linewidth | Shape Score |
| Baseline U-Net* | $0.02420 \pm 0.01711$ | $35.6 \pm 17.5$ | $0.172427 \pm 0.046778$ | $0.8253 \pm 0.1040$ |
| Deep Spectral Divers | **$0.01271 \pm 0.03592$** | **$44.2 \pm 10.6$** | $0.182649 \pm 0.030305$ | **$0.9486 \pm 0.2090$** |
| Spectralligence | $0.01697 \pm 0.02586$ | $33.9 \pm 13.9$ | **$0.166503 \pm 0.021886$** | $0.9195 \pm 0.2118$ |
| Dolphins | $0.07279 \pm 0.05466$ | $5.9 \pm 2.0$ | $0.191105 \pm 0.040382$ | $0.3700 \pm 0.4909$ |

### 3.1 Track 1: Simulated Data

Metric results for Track 1, simulated data, for each team are presented in Section 1 of Table 2. Representative reconstructed spectra for Track 1 are in Figure 1.

**Figure 1.**
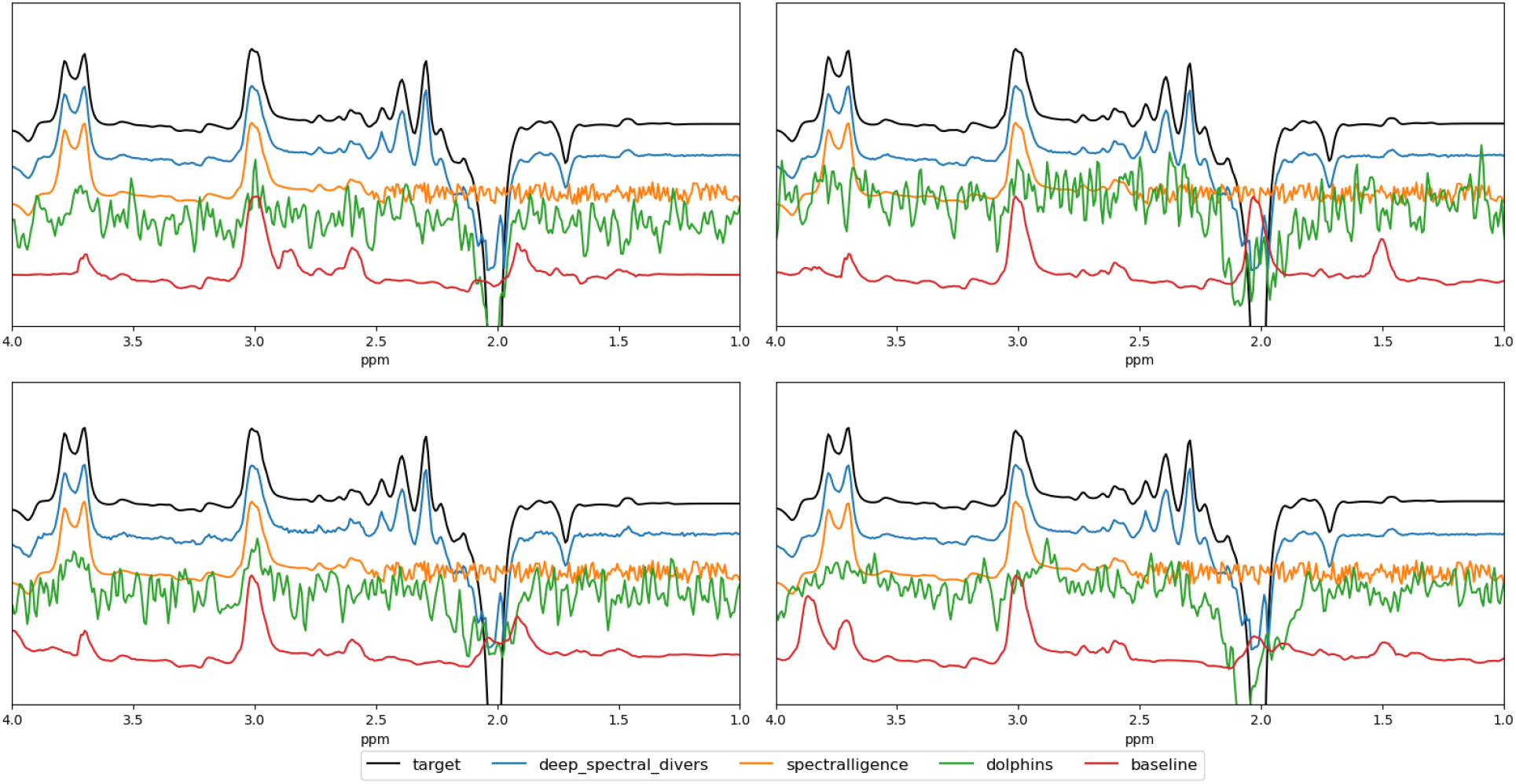
Four representative reconstructed final difference spectra for Track 1 - simulated data: U-Net baseline (red), Team Deep Spectral Divers (blue), Team Spectralligence (orange), Team Dolphins (green), and ground truth (black). Spectra are offset for better visibility. Both the blue and orange spectra are able to reproduce the GABA (3 ppm) and Glx (3.75 ppm) peaks well as compared to the target in black while the other DL reconstructions in red and green deviated considerably in one or both of the peaks.

### 3.2 Track 2: *In Vivo* Homogenous Data

Metric results for Track 2, homogeneous *in vivo* data, are presented in Section 2 of Table 1. Figure 2 shows four representative reconstructions by the different models compared to the target and Figure 3 is a box plot showing the distributions of the metrics among different models.

**Figure 2.**
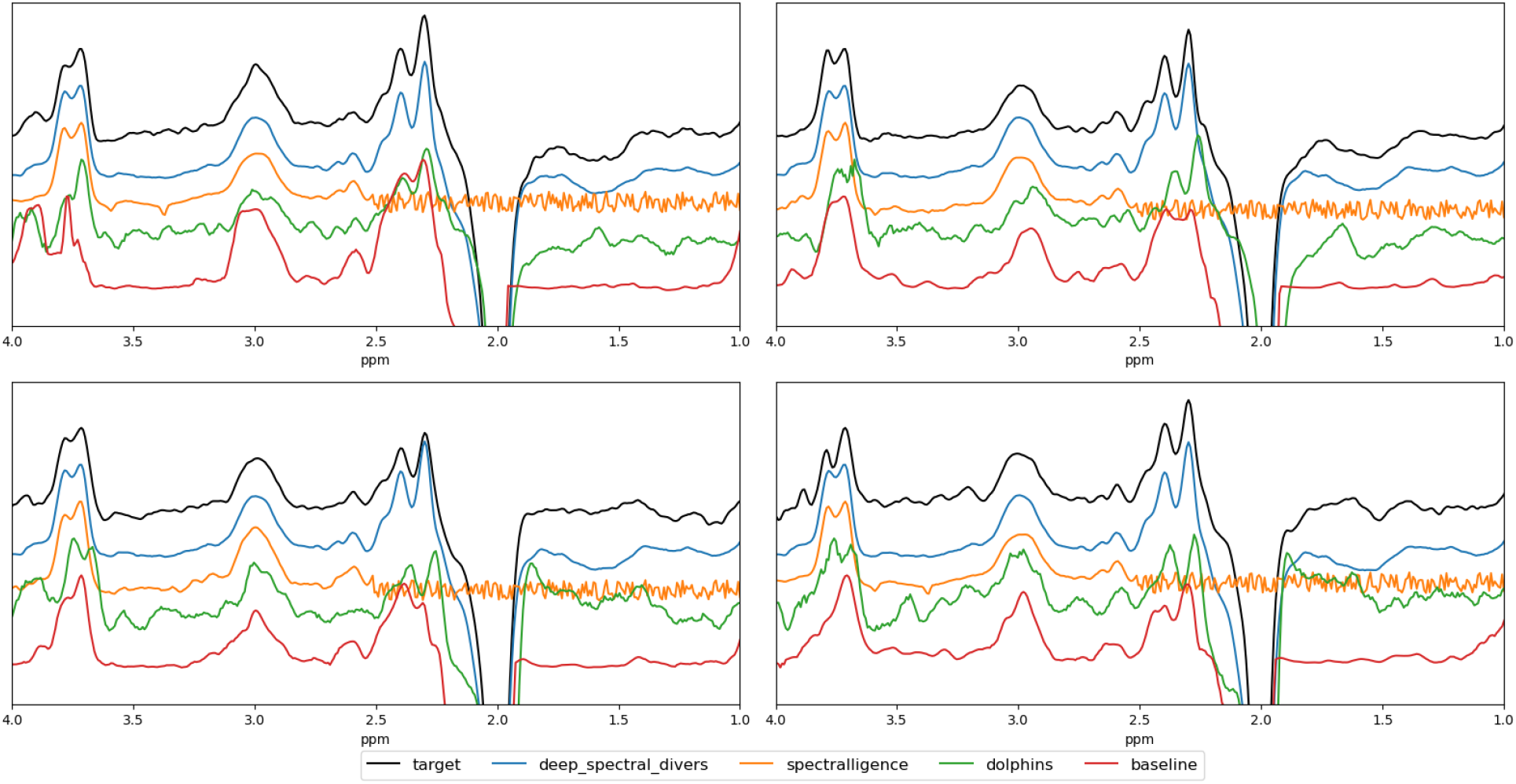
Four representative reconstructed final difference spectra for track 2 - homogeneous *in vivo* data: U-Net baseline (red), Team Deep Spectral Divers (blue), Team Spectralligence (orange), Team Dolphins (green), and target (full 320 transient reconstruction) (black). Spectra are offset for better visibility. All reconstructions follow the target spectra in black.The blue and orange spectra differ only in details of the peaks (such as subtraction artifacts or peak sharpness differences) while the red and green reconstructions present significant overall peak shape differences.

**Figure 3.**
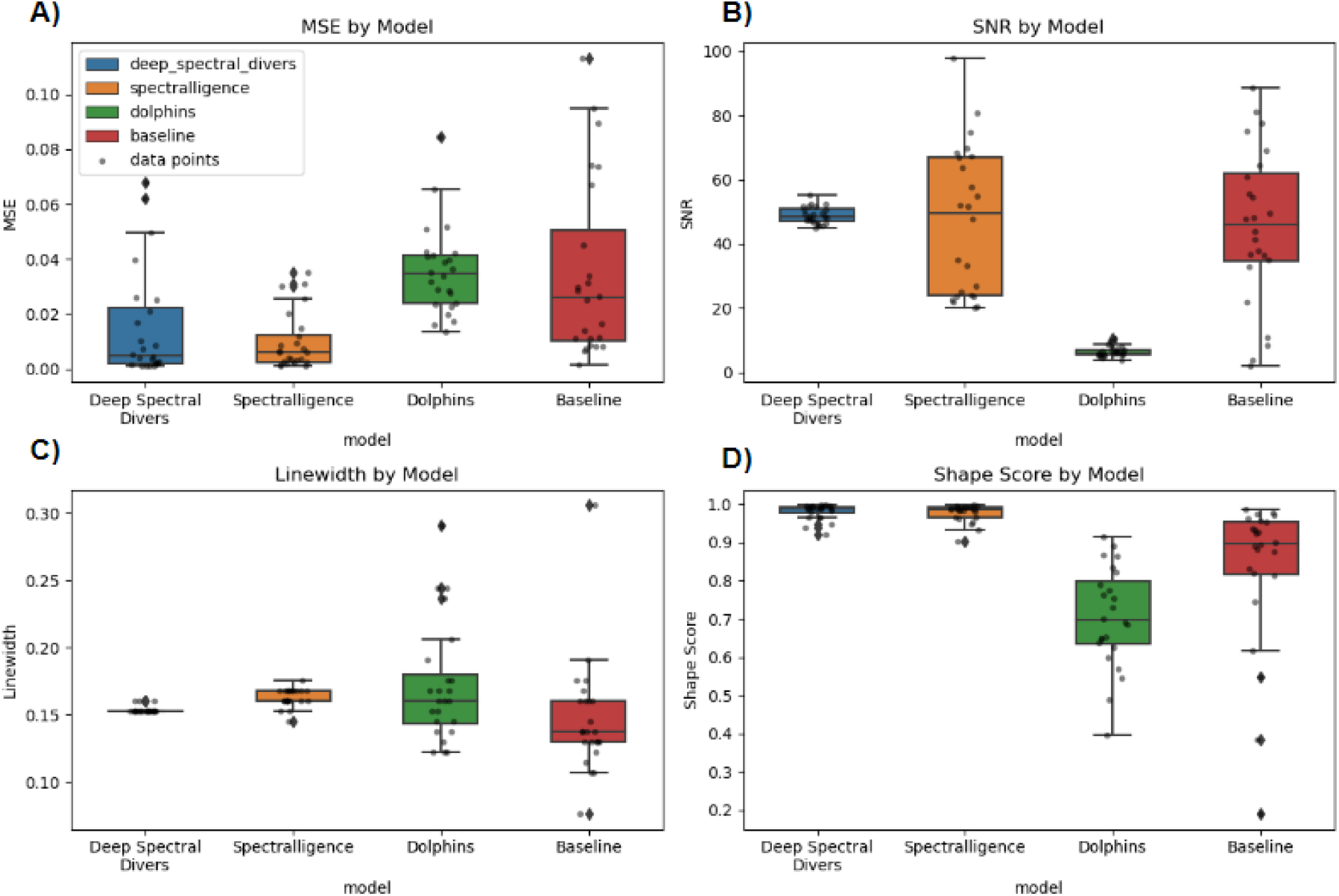
Box plots of the challenge metrics A) MSE, B) SNR, C) linewidth, and D) shape score for each team and the baseline for Track 2. Each team’s distribution is marked by a colored boxplot and each datapoint is marked by a gray dot. The color coding and order are the following: Deep Spectral Divers (Blue); Spectralligence (Orange); Dolphins (Green); and Baseline (red).

### 3.3 Track 3: *In Vivo* Heterogeneous Data

Metric results for Track 3, heterogeneous *in vivo* data, are presented in Section 3 of Table 2. Team Deep Spectral Divers obtained the best overall model similarly to Track 2. Figure 4 shows four examples of reconstruction by the different models compared to the target and Figure 5 is a box plot showing the means and distribution of each of the metrics among different models for track 3.

**Figure 4.**
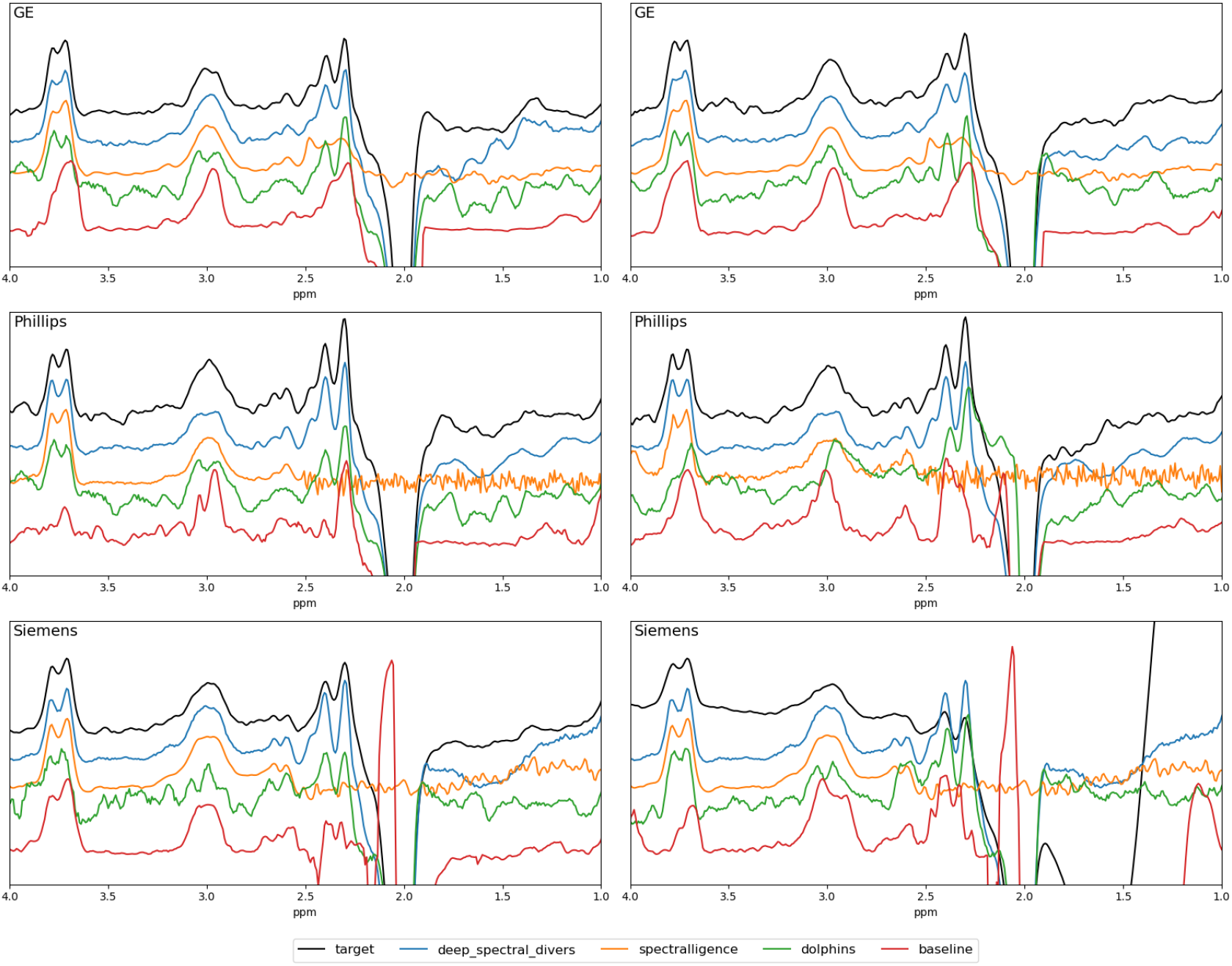
Sampled representative reconstructed final difference spectra for Track 3 - heterogeneous *in vivo* data from different vendors GE (top row), Philips (middle row), Siemens (bottom row): U-Net baseline (red), Team Deep Spectral Divers (blue), Team Spectralligence (orange), Team Dolphins (green), and target (full 320 transient reconstruction) (black). Spectra are offset for better visibility. GE DL reconstructions, in general, appear to follow the target better than Philips and Siemens likely due to having more training on GE scans from both Track 2 and 3. As seen in previous tracks, DL model outputs in blue and orange reconstruct key peak shapes (GABA and Glx) and details more precisely than those in green and red.

**Figure 5.**
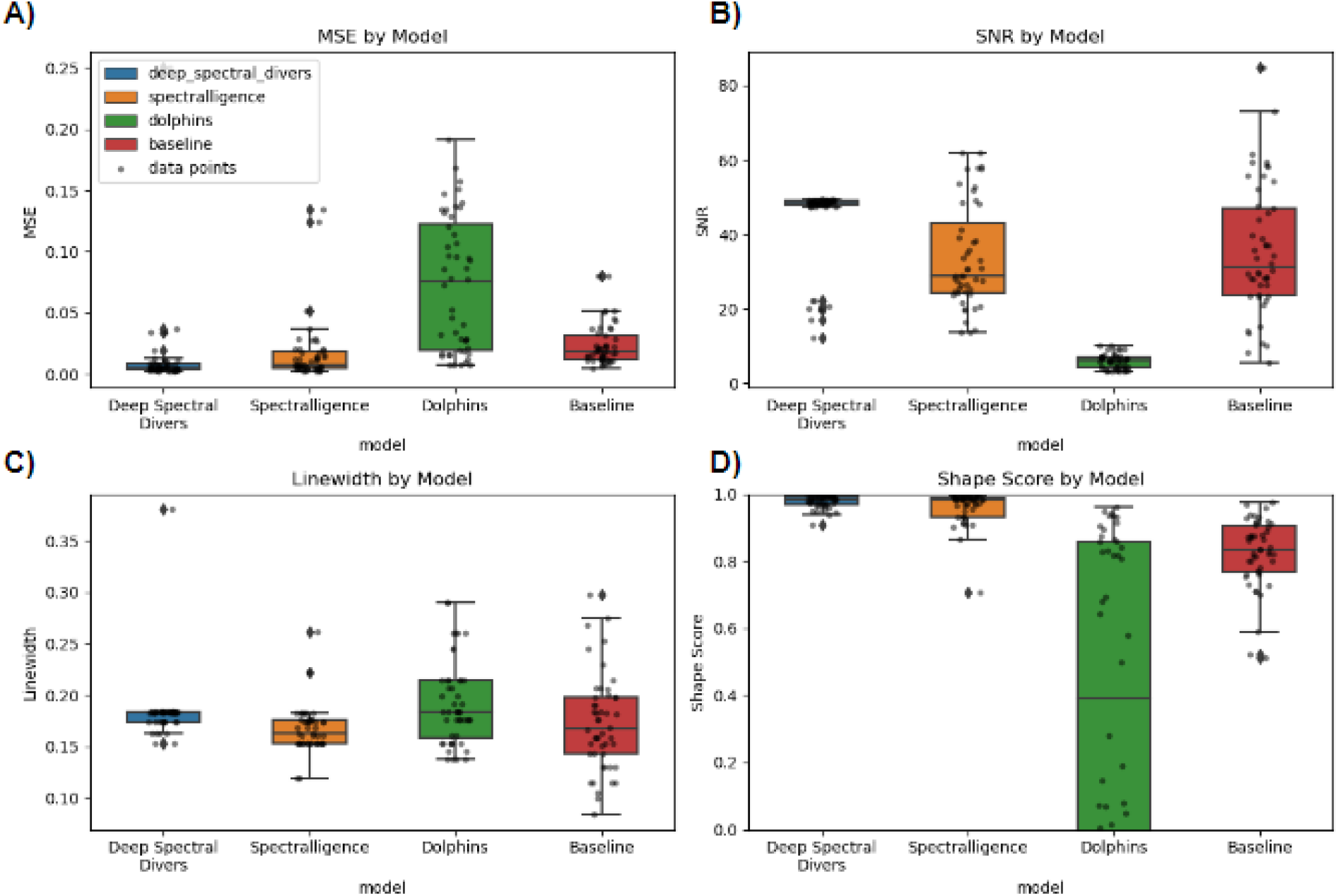
Box plots of the challenge metrics: A) mean squared error, B) SNR, C) linewidth, and D) shape score for each team and the baseline for Track 3. Each team’s distribution is marked by a colored boxplot and each datapoint is marked by a gray dot. The color coding and order are the following: Deep Spectral Divers (Blue); Spectralligence (Orange); Dolphins (Green); and Baseline (red).

#### 3.2.3 Results Validation: Fit Error

The fit error of reconstructions were obtained to validate *in vivo* results from Track 2 and Track 3. Fit errors from example reconstructed spectra from Track 2 and Track 3 are presented in Figure 6 and the distribution of the fit error for Track 2 and Track 3 for all submissions are shown in Figure 7.

**Figure 6.**
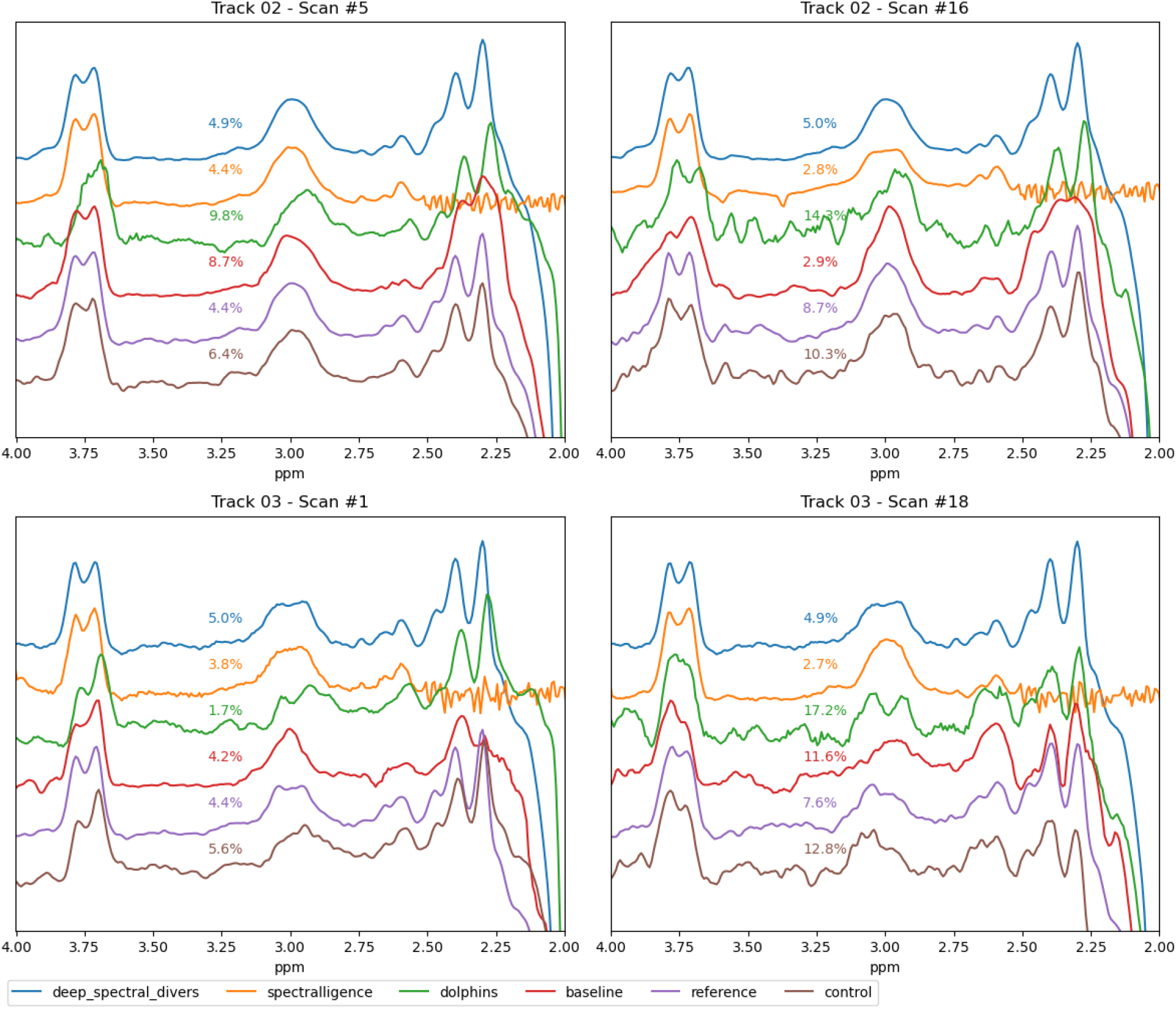
Representative sample reconstructed final difference spectra with associated fit error determined by Gannet for track 3 - heterogeneous *in vivo* data: U-Net baseline (red), Team Deep Spectral Divers (blue), Team Spectralligence (orange), Team Dolphins (green), reference (full 320 transient reconstruction) (purple), and control (first 80 transient reconstruction) (brown). Spectra are offset for better visibility. For Track 3, DL models that achieved the best results in the challenge, blue and orange, obtained more consistent fit errors across spectrum reconstructions than lower ranking challenge models, red and green.

**Figure 7.**
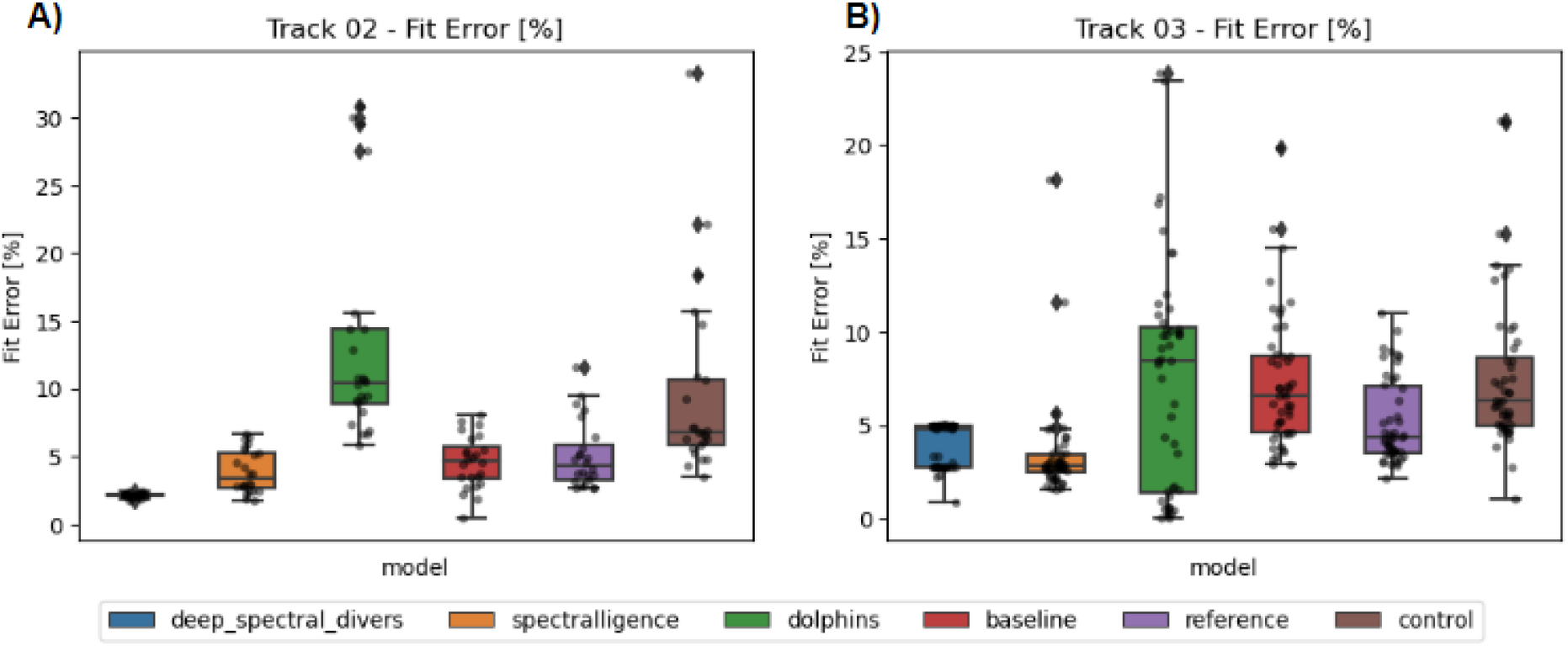
Box plots of fit errors for each team, the baseline, the reference, and control for Track 2 (A) and Track 3 (B). Each team’s distribution is marked by a colored boxplot and each datapoint is marked by a gray dot. The color coding and order are the following: Deep Spectral Divers (blue); Spectralligence (orange); Dolphins (green); Baseline (red); reference (purple); and control (brown). In general, mean fit error did not change remarkably for DL models from Track 2 which had homogeneous acquisition parameters to Track 3 which had heterogeneous acquisition parameters.

Since the fit error is a measure of reconstruction quantification fit, it is arguably closer to the objective of MR (*i.e.*, metabolite quantification) than the remaining metrics. Because of this, the correlations of fit error with the challenge metrics were also obtained. This may allow for the interpretation of the ability of the challenge metrics to predict not only spectral quality but also quantification accuracy. Figure 8 shows the relation between the challenge metrics and the fit error per scan per model along with the Pearson’s correlation coefficient. Correlations between fit error and MSE, linewidth, and shape score demonstrate a low degree of correlation while fit error and SNR demonstrate an inversely moderate degree of correlation.

**Figure 8.**
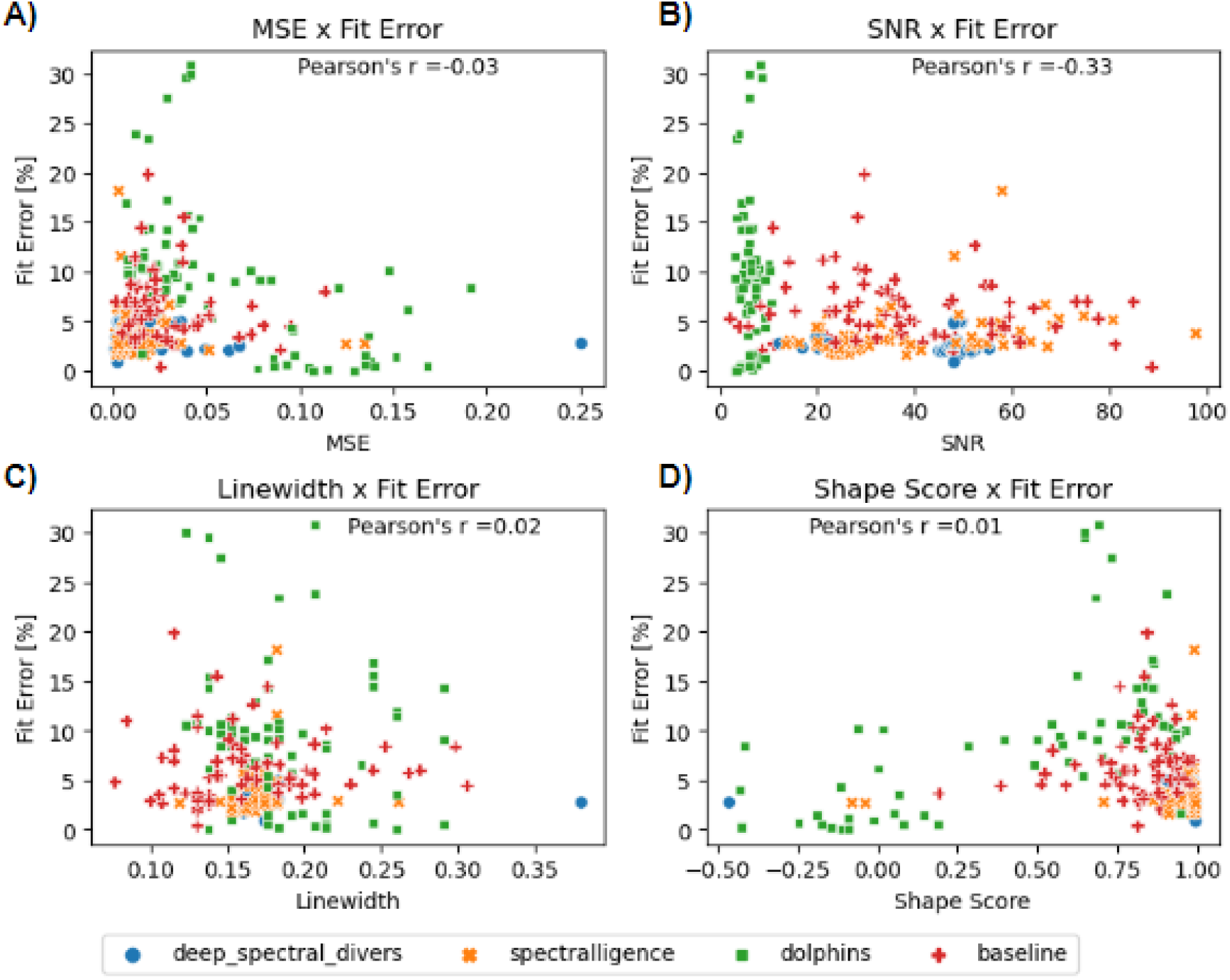
Distribution of correlation between fit error and challenge metrics for track 3. A) Correlation between fit error and mean squared error shown per transient, B) correlation between fit error and signal-to-noise ratio shown per transient, C) correlation between fit error and linewidth shown per transient, D), correlation between fit error and shape score shown per transient. Most models showed a small inverse correlation between fit error and SNR but lacked a correlation between fit error and the remaining evaluation metrics (MSE, linewidth, and shape score).

## 4 DISCUSSION

### 4.1 General Summary and Impact

Track 1 compared the reconstruction quality on simulated data between the baseline and all three team submissions, while Track 2 and Track 3 compared reconstruction quality on *in vivo* data. Models generally performed better on each metric for the simulated data in comparison to *in vivo* data. DL-reconstructed spectra were very similar to conventional reconstructions with 320 transients.

### 4.2 Model Architectures

While not an explicit objective of the challenge, participants had the opportunity to investigate different representations for spectroscopy data which is typically one dimensional by nature. In the challenge, all submissions used 2D inputs to the models after a transformation of the original 1D inputs. One team and the baseline used the concatenation of transients to form a 2D image, one team calculated a covariance matrix with the transients as the input, and one team constructed spectrograms from the transients.

It is interesting to see these transformations and the usefulness of turning the data from 1D to 2D on model selection, as it enabled teams to leverage the extensive DL literature on 2D representations and models.

### 4.2 Metric Assessment

Metrics for the challenge were chosen as a mix of conventional MRS and DL metrics, as well as a newly-proposed shape score metric. It included metrics that required a reference and self-contained metrics. The choices reflect the objective of analyzing different aspects of reconstructed spectra and staying close to existing literature.

MSE was selected as a metric that is more representative of DL quality as opposed to traditional MRS quality metrics (*i.e.* SNR and Linewidth). The MSE metric evaluated the error between 2.5 ppm and 4 ppm as this contains the GABA and co-edited Glx peaks while avoiding the direct effects of the editing pulse around 1.9 ppm. However, when assessing the spectrum as a whole for tracks 2 and 3, we note that although some models obtained a remarkably small MSE, reconstruction outside the window of interest did not have the expected metabolite peaks thus differing considerably from the target (full 320 transient reconstruction). It is hypothesized that this happens depending on model architecture due to the loss function not covering these regions. Although models which focus solely on optimizing quality within the window or region of interest may obtain better results for windowed metrics such as MSE, they may be sacrificing overall model reliability. In addition, while this metric does provide good insights into how the reconstruction relates to the full reconstruction, for *in vivo* data it cannot provide the objective assessment that a comparison to a ground truth can since even a preprocessed full reconstruction will contain data imperfections.

SNR is widely used in the field of MRS(Alexander Lin et al., 2021; In-Young Choi et al., 2021; Jamie Near et al., 2021). It has the limitation of only looking at the maximum signal of a single metabolite and an upfield region without peaks. For conventional MRS preprocessing, where we know that the noise throughout the entire spectrum is treated homogeneously, this is not an issue as the upfield region is an adequate proxy of the noise in the metabolites region. However, when considering DL black box models, that affirmation cannot be confirmed because a model could learn to decrease the amplitude of the signal (presumably noise) at that specific upfield region for SNR sampling in order to optimize the metric without improving the metabolite peak itself.

Linewidth is another metric widely used in the MRS field(Alexander Lin et al., 2021; In-Young Choi et al., 2021; Jamie Near et al., 2021) that is limited by the ambiguity of defining a threshold value. Intuitively, linewidth is set as a metric to minimize however with DL models it is possible to obtain results with linewidths smaller than the expected true value in which it becomes difficult to interpret whether a thinner or broader peak is better. Like SNR, this is a metric that becomes problematic when considering a DL black box, which if optimizing for this metric, could lead to a great improvement of the metric but may lead to unrealistic results. With the barriers to DL optimization of linewidth in combination with its poor predictive value of the top performing model in the challenge as seen by the baseline obtaining the best linewidth in two of the three tracks, we advise against using linewidth as an absolute value to evaluate future DL GABA-edited MRS reconstruction models.

The final challenge metric, shape score, requires a ground-truth like MSE, and so is limited in its application. As the metric was first proposed for the challenge, it is important to validate its results. When considering that the best submission for each track obtained the best ranking for this metric, we can conclude that the metric is likely a valid indicator of the quality of a reconstruction. Work to improve the scaling of this metric will be needed, especially for applications involving thresholding, as there is currently only a small difference in score between a subjectively good and bad reconstruction.

Given the challenge metrics, we see that each has its own limitations towards applicability in the opposite field (MRS vs. DL). For the MSE and shape score metrics traditionally made for DL applications, when applied to *in vivo* data, results cannot be adequately verified due to a lack of a ground truth. In contrast, for the SNR and linewidth metrics traditionally made for *in vivo* applications, when applied to DL-reconstructed data, results may be inadequate as models are optimizing the metrics in a way which is unrealistic. Changes to improve the robustness of metrics when used in a field they were not originally developed for may be needed. For instance, with linewidth, perhaps minimizing the absolute difference from the reference is more suitable for DL applications than minimizing the overall value. Nonetheless, the metrics complement each other as they look at different aspects of the reconstructions, allowing for a stronger qualitative assessment than each metric can provide individually.

### 4.3 Validation of Reconstruction

Fit error was evaluated as an additional metric with no influence on the ranking of submissions to determine the validation of the reconstructions. This metric was chosen as it is a widely accepted quantitative metric for spectrum quality assessment and provides a different perspective than the existing challenge metrics. Based on the challenge results seen in Figure 8, compared with other metrics used, fit error is only inversely moderately correlated with SNR. This correlation is not surprising, as quality is expected to improve with greater SNR and lower fit error. However, the lack of correlation with the other metrics indicates that they are complementary to the fit error in defining spectral quality.

An important note in the validation of the reconstruction is that all the metrics presented evaluate only a limited window or windows. For example, in figures 2-5, it is clear that team Spectralligence obtained good metric results. Nonetheless, the signal under 2.5 ppm, outside the region of interest, is very different from what is expected. This raises the question if a metric that evaluates the whole spectrum should also be used or if regions such as these should be discarded when working with DL-based reconstructions, as they are not relevant for quantification itself.

## 5. CONCLUSION

A challenge was proposed to establish a quality benchmark for deep learning GABA-edited MRS reconstructions using four times less data (80 transients) than a full data reconstruction (320 transients). Track 1 used simulated data, track 2 used *in vivo* data with homogeneous parameters and track 3 used *in vivo* data with heterogeneous parameters. While teams only had two months to prepare submissions, the results demonstrated that high quality reconstructions will achieve favorable results across most metrics with the best solution of each track obtaining the best results in three of the four metrics. As anticipated, simulated data results for all teams were generally better than *in vivo* results suggesting more work is needed when fine tuning (generalizing) models from simulated to *in vivo* data. In addition, metrics used for applications which they were not intended for (*in vivo* MRS vs. DL) should be done cautiously as each individual metric presents its own limitations which were, for the most part, overcome by the combination of all metrics.

## DATA AND CODE AVAILABILITY

All code pertaining to the simulated data generation or deep learning models in this article can be found at the following github repository: https://github.com/rmsouza01/Edited-MRS-challenge. *In vivo* data is available in Big GABA public repository (Mikkelsen et al., 2017).

## AUTHOR CONTRIBUTIONS

RB, HB, RS, and ADH were responsible for designing and organizing the challenge, providing baseline data and code, tutorials on how to use these, results presentation, and creating and maintaining a challenge website and code repository. These authors were also responsible for writing the majority of this manuscript. GD, MO, LU, SD, PDPC, and LR were responsible for the Deep Spectral Divers submission. JPM, DMJVDS, SA, GSD, MV, JFAJ, MB, and RJGVS were responsible for the Spectralligence submission. AQ, CR, and SN were responsible for the Dolphins submission. All authors provided feedback throughout the writing process and reviewed and approved the final submitted version.

## ETHICS STATEMENT

The work completed in this manuscript adheres to the ethical guidelines outlined by MIT Press.

